# Analysis of genetic diversity and association of seed and mucilage yields with inter simple sequence repeats (ISSR) polymorphism in a collection of *Plantgao* species

**DOI:** 10.1101/2020.07.06.189266

**Authors:** Bagheri Motahareh, Bahram Heidrai, Zolfaghar Shahriari, Ali Dadkhodaie, Zahra Heidari, Christopher M Richards

## Abstract

Analysis of genetic diversity in medicinal plants assists germplasm conservation and selection for use in breeding schemes. The aims of the present study were to assess genetic diversity and differentiation of several *Plantago* species using Inter Simple Sequence Repeats (ISSR) markers and identify marker-trait associations (MTAs). Thirty-one *Plantago* accessions belonging to eight species with various mating system and chromosome number were collected from geographical regions of Iran environments. Polymorphism in the DNA of *Plantago* accessions were analyzed using polymerase chain reaction (PCR) of 25 ISSR primers. The data for number of polymorphic bands were analyzed on the basis of several genetic diversity parameters. The results of gel analysis indicated that the ISSR primers amplified 5 to 21 polymorphic bands with 100 to 3000 bp size. The mean polymorphism was 83.83% and five primers showed 100% polymorphism among *Plantago* accessions. The Polymorphism Information Content (PIC) for ISSR as a dominant marker ranged from 0.1103 to 0.3829 with the mean 0.2727 in the species tested. Accessions in *P. amplexicaulis* and *P. pysillum* species represented the highest Nei’s and Shannon’s genetic diversity whilst the lowest obtained for *P. lagopus*. Analysis of phylogenetic network generated by the Neighbor-Net Algorithm showed moderate split of the eight species tested and the network depicted moderate conflict. The principal coordinate analysis (PCoA) results showed lower conflict in separation of accessions of the eight species. Fifty-six significant MTAs were detected for the traits tested in *Plantago* accessions, of which six were shared between three seed and mucilage traits and 24 were common between two traits. The coefficient of determination (R^2^) for the identified MTAs varied between 32 and 73%. In conclusion, the results of genetic diversity analysis suggested that ISSR marker could efficiently differentiate *Plantago* species and the information of genetic diversity might assist *Plantago* improvement and conservation.

## Introduction

*Plantgao* which is the most prevalent genera of the family Plantaginaceae comprises of over 400 species many of which are cosmopolitan weeds [1]. They are found all over the world in many different habitats, most commonly in wet areas like seepages or bogs. Plants in the genus *Plantago* are growing to 60 cm tall with annual and perennial growth habits and are found in a wide range of habitats including deserts, sea cliffs and tropical mountains [1,2]. Some of *Plantago* species are native of Persia and some species are widely cultivated in India, Pakistan and Mexico [2,4]. Seed of *Plantago* possesses industrial and medical uses. Mucilage is a thick, gluey and polysaccharide substance [2,5,6]. Seed mucilage has various uses comprising paper production, gum and medicine industries [2]. Despite its industrial and medicinal uses, the productivity of *Plantago* species is poorly understood. A first step in utilization is to determine the genetic identity and relatedness among accessions within the collection [1,2].

The International Union for Conservation of Nature has highlighted the importance of conservation of *Plantago* species [7]. Genetic diversity is the basis of plant conservation, environmental survival and adaptation [8]. Identification of the genetic diversity of endangered species for preserving germplasm is one of the main goals of conservation strategies [8,9]. Species delineation helps the preservation of within and among population variations for long-term uses [8]. Markers in DNA level are efficient tools to dissect and identify genetic variation in plant populations [10,11].

Inter simple sequence repeats (ISSR) tool has been widely used in phylogenetic and genetic variation analysis of plant species [12,13,14,15,16,17]. Inter simple sequence repeats markers were used for analysis of genetic structure and population differentiation in *Plantago brutia* [18]. Ferreira et al. [9] assessed the ability of ISSR markers for detection of genetic diversity of *P. almogravensis* and *P. algarbiensis* populations and demonstrated that information of ISSR can be used to for rational conservation decisions. Recently, Osman and Abedin [1] used five ISSR primers to analyze five species of *Plantago* collected from different regions of Saudi Arabia.

Identification of polymorphic DNA markers linked to seed yield might assist construction of genetic maps and marker- assisted (MAS) selection programs for development of new varieties in the non-model plant species. Although the diversity of *Plantago* species has been assessed in previous studies at morphological and DNA levels, few species have been tested in most of studies and less efforts have been devoted to analyze marker-trait associations in this species. The aims of the present study were to (1) evaluate collection diversity of *Plantago* to identify variations in accessions belonging to various species and (2) analysis of association of ISSR markers with seed and mucilage traits. Such information will contribute to the description of ex situ diversity and assist in pre breeding applications.

## Materials and Methods

### Plant Material and field evaluations

The seeds of 31 accessions belonging to eight species of *Plantago* were collected from different parts of Iran environments with various eclogical characteristics. A bulk of seeds of several plants were collected for each species in a confined area of various Iran regions. A map showing locations for sampling *Plantago* accessions is shown in Figure 1. Metorology and geographical chcracteristics of the collection areas and accession codes are listed in Table 1. The growth, mating system and chromosome charactertstics of the species tested are presented in Table 2.

**Figure 1.**
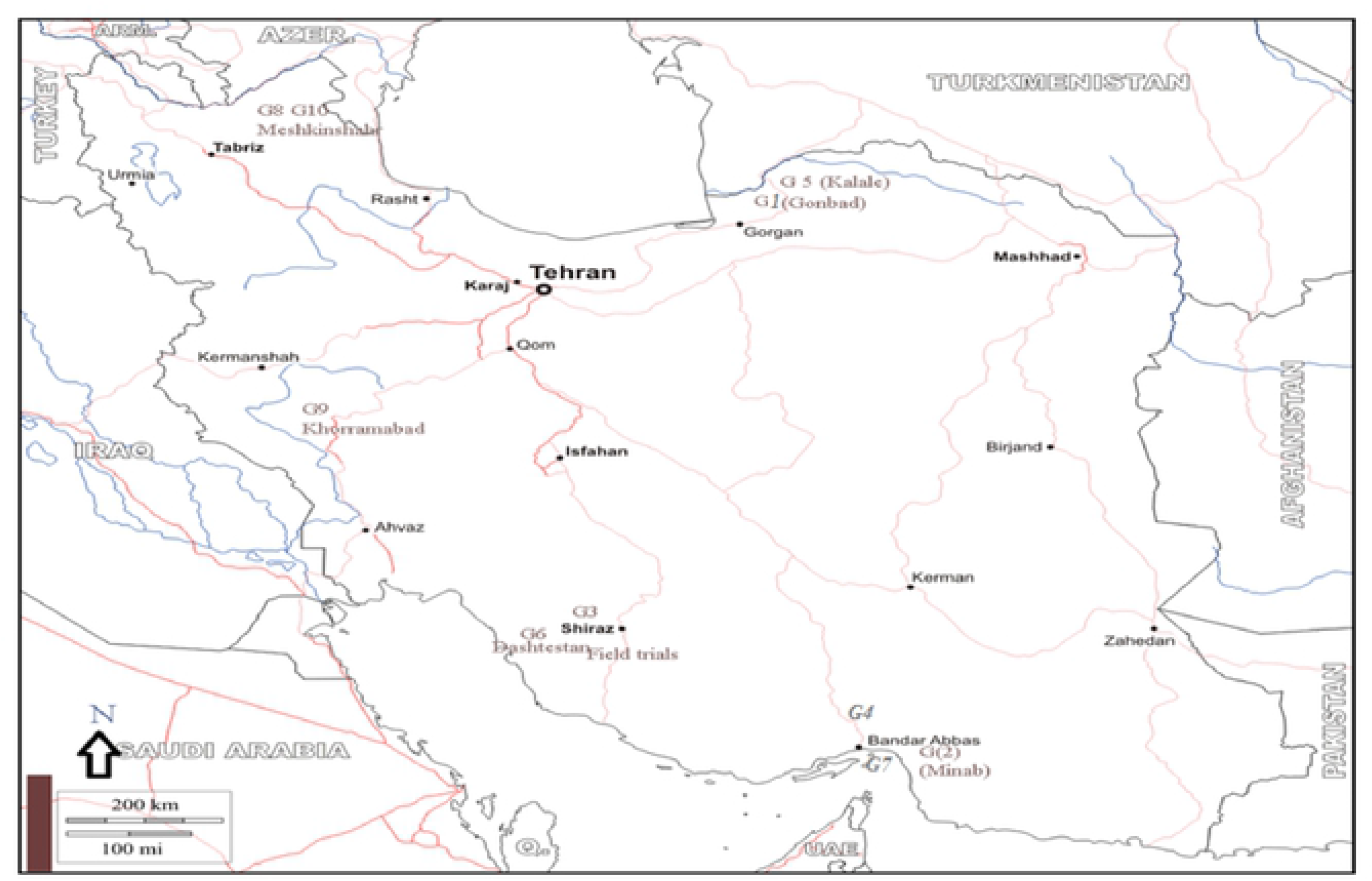
The Iran map showing places where the *Plantago* species were collected. The G letter and numbers refer to the name of species in Table 1.

**Table 1.** Meteorological and geographical data for Iran areas where *Plantago* species were collected

| Accession<br>code | Species | Province/City | Low<br>Temp*<br>(°C) | High<br>Temp<br>(°C) | Sea<br>level<br>height<br>(m) | Latitude | Longitude | Mean<br>Temp<br>(°C) | Annual<br>rainfall<br>(mm) | Climate<br>classification |
| --- | --- | --- | --- | --- | --- | --- | --- | --- | --- | --- |
| G1 | *P. major* | Shahzand/Markazi | -3.3 | 24.6 | 1906 | 33.56 | 49.24 | 11.3 | 312 | Dsa |
| G2 | *P. major* | Ardebil/Ardebil | -0.2 | 21.9 | 1391 | 38.14 | 48.18 | 12.1 | 381 | Csb |
| G3 | *P. major* | Sanandaj/Kordestan | -0.4 | 26.2 | 1499 | 32.25 | 47 | 12.8 | 492 | Csa |
| G4 | *P. major* | Shahrood/Semnan | 1.7 | 25.7 | 1351 | 28.01 | 56.21 | 14.2 | 162 | Bwk |
| G5 | *P. major* | Najaf abad/Esfahan | 1.9 | 27.5 | 1650 | 32.49 | 51.49 | 15 | 155 | Bsk |
| G6 | *P. major* | Behshahr/Mazandaran | 8.5 | 26.8 | 16 | 38.01 | 56.19 | 17.3 | 522 | Csa |
| G7 | *P. major* | Shiraz/Fars | 6 | 28.1 | 1536 | 29.45 | 53 | 16.8 | 316 | Bsk |
| G8 | *P. officinalis* | Gorgan/Golestan | 8.2 | 27.8 | 133 | 37 | 54.3 | 17.8 | 515 | Csa |
| G9 | *P. officinalis* | Khorram abad/Lorestan | 5 | 29.6 | 1185 | 33.3 | 48.3 | 16.9 | 488 | Csa |
| G10 | *P. officinalis* | Arak/Markazi | -5.5 | 26.1 | 1752 | 34.25 | 49.35 | 11.8 | 316 | Dsa |
| G11 | *P. amplexicaulis* | Pars abad/Ardebil | 0.7 | 21.9 | 1391 | 39.39 | 47.54 | 12.1 | 382 | Bsk |
| G12 | *P. ovata* | Gonbad/Golestan | 8.3 | 28.1 | 164 | 38.3 | 44.54 | 17.7 | 435 | Bsk |
| G13 | *P. ovata* | Minab/Hormozgan | 18.4 | 34.8 | 42 | 27.01 | 56.54 | 27.3 | 123 | Bwh |
| G14 | *P. ovata* | Kalale/Golestan | 7.3 | 28.5 | 155 | 37.38 | 55.3 | 17.4 | 315 | Bsk |
| G15 | *P. ovata* | Shiraz/Fars | 6 | 28.1 | 1536 | 29.45 | 53 | 16.8 | 316 | Bsk |
| G16 | *P. ovata* | Sirmand/Hormozgan | 11.5 | 31 | 928 | 27.59 | 56.07 | 21.7 | 147 | Bwh |
| G17 | *P. pysillum* | Dashtestan/Booshehr | 14.3 | 32.6 | 66 | 29.3 | 50.45 | 24.2 | 216 | Bwh |
| G18 | *P. pysillum* | Khoram abad/Lorestan | 5 | 29.6 | 1185 | 33.3 | 48 | 16.9 | 488 | Csa |
| G19 | *P. pysillum* | Bandar abbas/Hormozgan | 18.3 | 34.3 | 8 | 27.45 | 56 | 27.2 | 136 | Bwh |
| G20 | *P. pysillum* | Meshkin shahr/Ardebil | -1.7 | 21.4 | 1453 | 38.3 | 47.5 | 9.7 | 356 | Csb |
| G21 | *P. pysillum* | Ardebil/Ardebil | -1.7 | 21.4 | 1453 | 38.3 | 47.5 | 9.7 | 356 | Csb |
| G22 | *P. lanceolata* | Salmas/East Azarbajejan | -2.4 | 23.3 | 1378 | 38.11 | 44.46 | 10.5 | 388 | Csa |
| G23 | *P. lanceolata* | Estahban/Fars | 5.7 | 28.2 | 1736 | 29.07 | 54.04 | 17 | 222 | Bsk |
| G24 | *P. lanceolata* | Arak/Markazi | -5.5 | 26.1 | 1752 | 34.25 | 49.35 | 11.8 | 316 | Dsa |
| G25 | *P. lanceolata* | Semirom/Esfahan | -1.8 | 24.1 | 2372 | 31.25 | 51.34 | 11.6 | 235 | Csa |
| G26 | *P. lanceolata* | Andimeshk/Khozestan | 11.7 | 36.5 | 145 | 32.31 | 48.22 | 24.2 | 358 | Bsh |
| G27 | *P. coronopus* | Agh ghala/Golestan | 8.3 | 28.4 | -14 | 36.55 | 54.2 | 18.1 | 432 | Bsh |
| G28 | *P. coronopus* | Haji abad/Hormozgan | 11.5 | 31 | 928 | 27.18 | 55.54 | 21.7 | 147 | Bwh |
| G29 | *P. lagopus* | Dashtestan/Booshehr | 14 | 34.8 | 66 | 29.3 | 50.45 | 24.7 | 211 | Bwh |
| G30 | *P. lagopus* | Dehloran/Ilam | 9.7 | 34.4 | 220 | 32.41 | 47.15 | 22.5 | 280 | Bsh |
| G31 | *P. amplexicaulis* | Dashtestan/Booshehr | 14 | 34.8 | 66 | 29.3 | 50.45 | 24.7 | 211 | Bwh |

Temp.: temperature

**Table 2.**
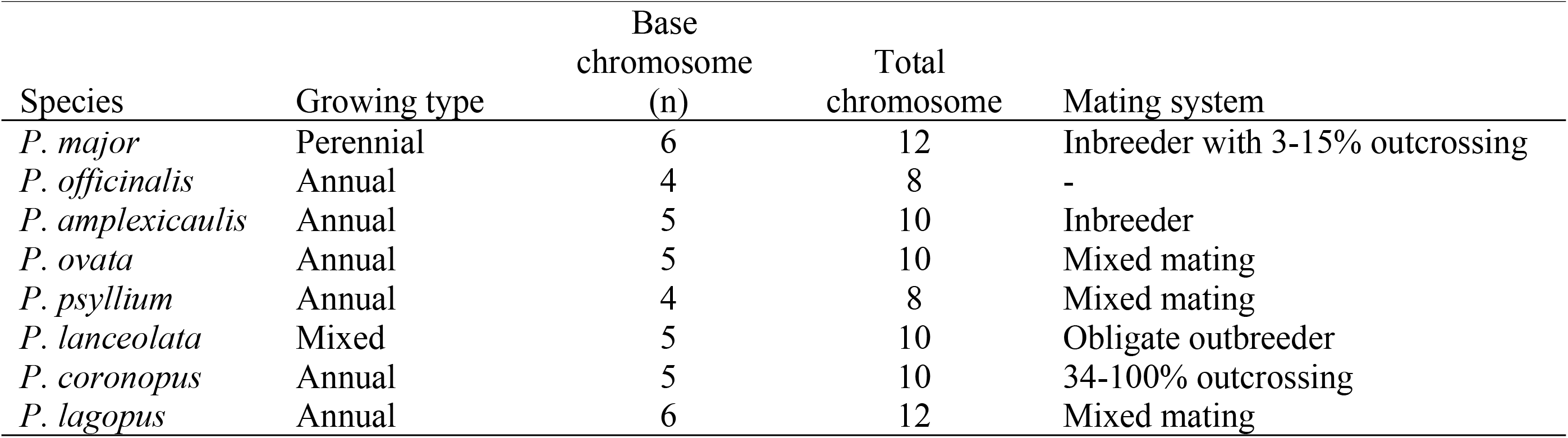
Growth, mating system and chromosome characteristics of *Plantago* species tested for genetic diversity.

| Species | Growing type | Base<br>chromosome<br>(n) | Total<br>chromosome | Mating system |
| --- | --- | --- | --- | --- |
| *P. major* | Perennial | 6 | 12 | Inbreeder with 3-15% outcrossing |
| *P. officinalis* | Annual | 4 | 8 | - |
| *P. amplexicaulis* | Annual | 5 | 10 | Inbreeder |
| *P. ovata* | Annual | 5 | 10 | Mixed mating |
| *P. psyllium* | Annual | 4 | 8 | Mixed mating |
| *P. lanceolata* | Mixed | 5 | 10 | Obligate outbreeder |
| *P. coronopus* | Annual | 5 | 10 | 34-100% outcrossing |
| *P. lagopus* | Annual | 6 | 12 | Mixed mating |

Managed field trials were performed in two locations, Shiraz and Marvdasht. In Shiraz, *Plantago* was grown under two irrigation regimes (drought stress and normal irrigation) in three consecutive years (2013-2014-2015) at the Research Farm of School of Agriculture (52° 46’N, 29° 50’ E). A two-year (2014-2015) experiment under two irrigation regimes was performed at Kooshkak Agricultural Research Station (52° 34’N, 30° 7’ E), Marvdasht, Iran. The experimental layout in the managed trials in both locations was a randomized complete block design (RCBD) with three replicates. For normal irrigation regime, irrigation was performed throughout growing season while for the drought stress treatment water deficit treatment was imposed as 50% field capacity (FC) started at the 2-3 leaf stage of growth. A seed density of 4 kg/ha was used for sowing in the 1.2 × 1 m experimental plots with two rows 60 cm apart. Seeds were planted in early April in managed field trials in both Shiraz and Kooshkak. 30 kg N ha^−1^ and 30 kg P_2_O_5_ ha^−1^ were added to the soil at sowing. Seed yield (g/plant), mucilage yield (g/plant) and mucilage content (as percentage per 100 g seed) were measured for each genotype in each trial. Mucilage content was extracted based on a method explained by Sharma and Koul (1986). Mucilage yield was calculated as g/plant.

### DNA Extraction and ISSR Analysis

The seeds were sown in 5 kg pot and fresh leavesof developed seedlings were used for DNA extcarion. DNA samples from young leaves were extracted using CTAB method [19]. DNA concentration and quality was measured using a spectrophotometer (U-1800 Hitachi, Japan) and agarose gel electrophoresis. The DNA was diluted to a concentration of 10 ng mL^−1^. Twenty-five ISSR primers were used for assessment of genetic diversity in the *Plantago* populations (Table 3). Polymorphism chain reaction (PCR) was performed in a reaction volume of 20.0 μL containing 1.0 μL genomic DNA, 2.0 mL primer, 5.0 μL master mix (10 mM) and 12.0 μL water. The ISSR amplification program was run as outlined in (Suppl. Table 1). DNA amplicons were differentiated in 2% agarose gel at 80 W for 6 hours in 1X TBE buffer (100 mM Tris-Borate, pH 8.0, 2 mM EDTA). Total and polymorphic bands and band size were determined using GelAnalyser software (Suppl Fig. 1).

**Table 3.**
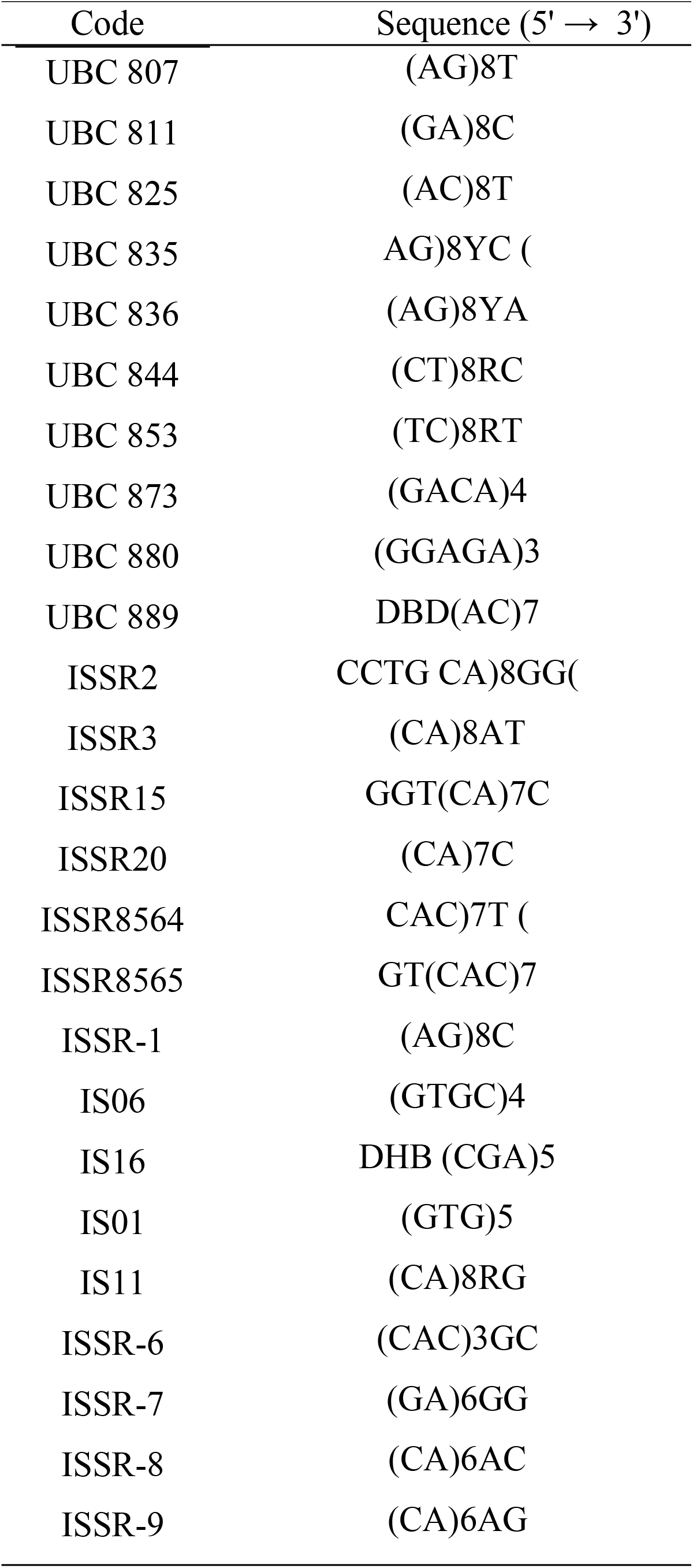
Sequence for Inter simple sequence repeats primers used for genetic diversity analysis in *Plantago* species

### Data analysis

Amplified fragments were scored using GelAnalyser 2010 (Suppl. Fig. 1). The binary data of ISSR markers were analyzed using POPGENE version 32 [20]. Shannon’s information index [21] was estimated as follows:

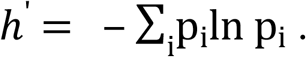

where pi is the frequency of allele in a locus

Polymorphic Information Content (PIC) was calculated followed by an equation proposed for dominant markers [22],

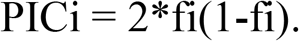

fi is the frequency of the amplified allele (present) and (1-fi) is the frequency of the absent allele (missing).

Marker index (MI) was calculated for each ISSR primer using an equation as below [23]:

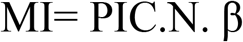

where β is the fraction of polymorphic markers, estimated after considering the polymorphic loci (np) and non-polymorphic loci (nnp) as β = np/(np+ nnp).

Coefficients of Nei’s genetic distance were calculated followed by Nei [24,25].

Nei’s genetic distance (D) was based on the identity of genes between two populations that formulated as,

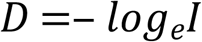

where, *I* is the normalized identity of genes between two populations [24]. Genetic diversity parameters (*h*, *I* and polymorphism) were normalized based on linear adjustment of allelic richness by unequal number of accessions per species tested [26].

The structure of *Plantago* species was assessed using principal component analysis (PCA) followed by Neighbour-Joining clustering methods. A reticulate network was constructed using the NeighborNet algorithm [27] that was implemented in SplitsTree 4.14.4 [28]. Phylogenetic networks are generalizations of phylogenetic trees that represent reticulate signals. The network displays relative evolutionary distances between taxa as well as uncertainty in the groupings in the form of “splits” (or “webbing”) of internal branches.

### Analysis of MTA

Analysis of association of the ISSR markers and seed and mucilage traits was performed using TASSEL version 4.2.1 [29]. The mean data for each accession was used to identify MTAs. The GLM was used to obtain the Q-matrix at maximum ΔK. The kinship matrix (K-matrix) was calculated based on the results of marker data in TASSEL software [9. In the GLM model, no control was imposed on population structure and relative kinship values [30]. The means of traits (P-matrix) were used to identify stable MTAs. In TASSEL software, a Manhattan plot showing Log10 p-values for ISSR markers was used to remove spurious associations and identify highly significant MTAs with prominent contribution to phenotype variation [31,32].

## 3. Results

### Variation in amplified bands and polymorphism level

The gel image for the UBC825 and UBC836 primers showing variation in the amplified bands in *Plantago* species is presented in Fig. 2 and 3. Number of bands amplified by ISSR primers ranged from five for ISSR-7 to 21 for ISSR-9 (Table 4). Allele size for ISSR primers varied between 100 and 3000 bp. The largest variation was 200-3000 bp belonging to the amplicons of the ISSR-8 (Table 4). The mean polymorphism was 83.83% and five primers represented 100% polymorphism with respect to the bands amplified in *Plantago* accessions. Except eight primers, the reminders produced higher than 10 polymorphic bands in the *Plantago* genotypes. The PCR for the ISSR-9 primer resulted in the highest polymorphic bands (19 bands) whilst three bands were identified for the UBC853 primer. The range for PIC was between 0.1103 and 0.3829 with the mean 0.2727 in the species tested. The ISSR9 and IS01 primers showed high PIC in the species tested. Marker index varied between *Plantago* species with mean 3.38. The UBC853 and ISSR-9 primers showed the lowest (0.37) and highest (7.28) marker index, respectively.

**Figure 2.**
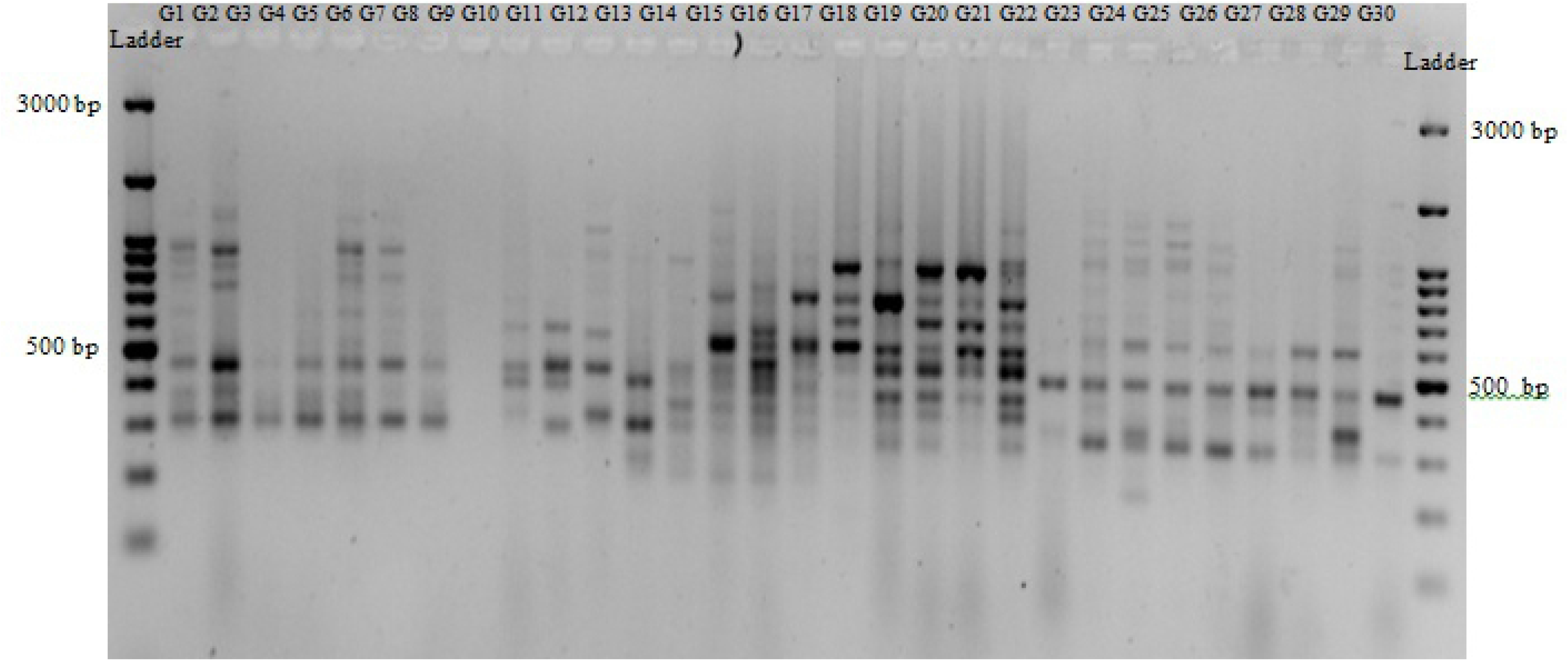
The gel image for the UBC825 primer showing variation in the amplified bands in *Plantago* species

**Figure 3.**
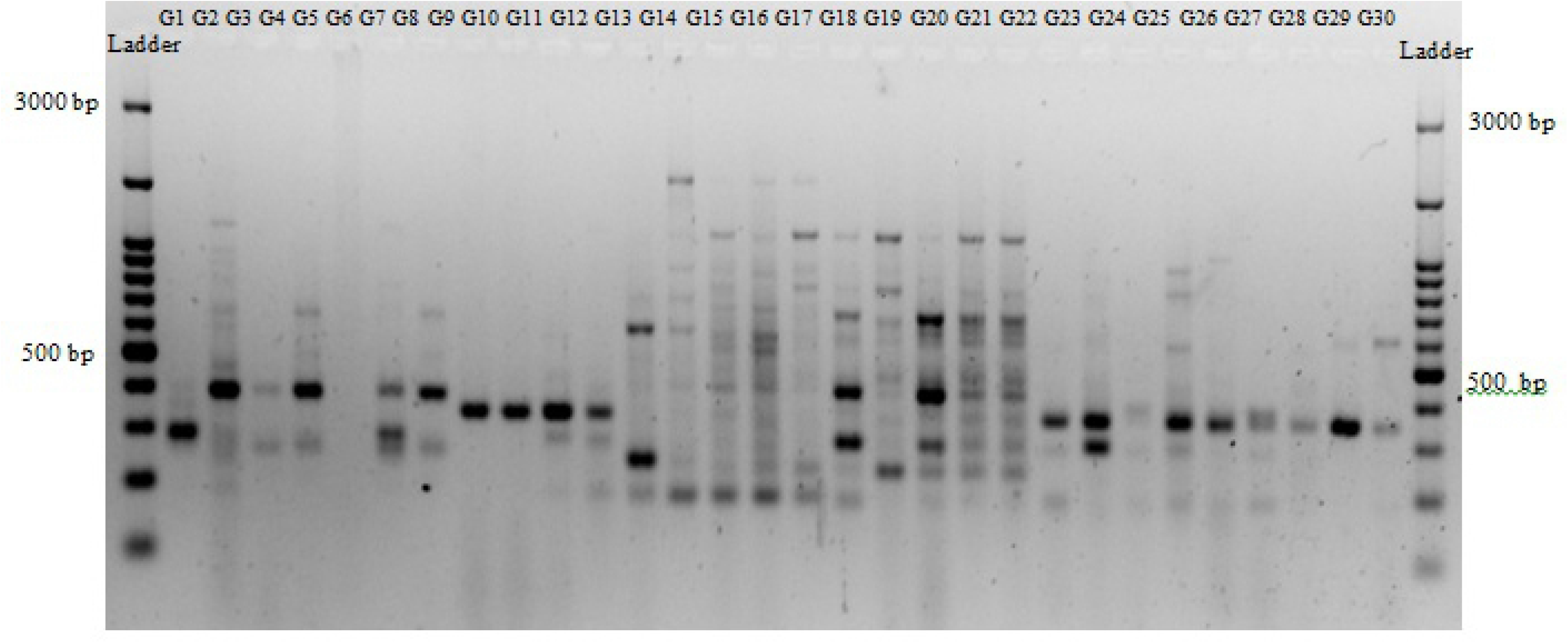
The gel image for the UBC836 primer showing variation in the amplified bands in *Plantago* species

**Table 4.** Polymorphism indices for inter simple sequence repeats (ISSSR) primers used for genetic diversity analysis in *Plantago*

| Primer codes | Total band number | Band size (bp) | Polymorphic band | $\beta$: Polymorphism (%) | PIC | Marker Index |
| --- | --- | --- | --- | --- | --- | --- |
| UBC807 | 15 | 150-2250 | 13 | 86.67 | 0.2788 | 3.62 |
| UBC811 | 18 | 140-2500 | 17 | 94.44 | 0.3388 | 5.76 |
| UBC825 | 17 | 290-1500 | 16 | 94.12 | 0.3560 | 5.70 |
| UBC835 | 9 | 180-1000 | 8 | 88.89 | 0.2516 | 2.01 |
| UBC836 | 17 | 200-1800 | 14 | 82.35 | 0.3064 | 4.29 |
| UBC844 | 13 | 140-980 | 8 | 61.54 | 0.2388 | 1.91 |
| UBC853 | 7 | 340-900 | 3 | 42.86 | 0.1219 | 0.37 |
| UBC873 | 11 | 220-1500 | 8 | 72.73 | 0.2887 | 2.31 |
| UBC880 | 11 | 240-2000 | 5 | 45.45 | 0.1324 | 0.66 |
| UBC889 | 11 | 200-1000 | 10 | 90.91 | 0.3012 | 3.01 |
| ISSR2 | 6 | 250-950 | 4 | 66.67 | 0.1103 | 0.44 |
| ISSR3 | 13 | 200-1000 | 11 | 84.62 | 0.2494 | 2.74 |
| ISSR15 | 18 | 250-950 | 14 | 77.78 | 0.2509 | 3.51 |
| ISSR20 | 13 | 220-800 | 13 | 100.00 | 0.2916 | 3.79 |
| ISSR8564 | 8 | 220-1500 | 6 | 75.00 | 0.1368 | 0.82 |
| ISSR8565 | 19 | 200-1200 | 15 | 78.95 | 0.2541 | 3.81 |
| ISSR-1 | 10 | 100-650 | 10 | 100.00 | 0.3329 | 3.33 |
| IS06 | 17 | 210-1400 | 15 | 82.24 | 0.2297 | 3.22 |
| IS16 | 15 | 190-1600 | 15 | 100.00 | 0.3464 | 5.20 |
| IS01 | 16 | 170-750 | 16 | 100.00 | 0.3816 | 6.11 |
| IS11 | 12 | 200-750 | 12 | 100.00 | 0.3201 | 3.84 |
| ISSR-6 | 15 | 200-1800 | 15 | 100.00 | 0.3660 | 5.49 |
| ISSR-7 | 5 | 150-1200 | 4 | 80.00 | 0.2622 | 1.05 |
| ISSR-8 | 15 | 200-3000 | 15 | 100.00 | 0.2872 | 4.31 |
| ISSR-9 | 21 | 200-1500 | 19 | 90.48 | 0.3829 | 7.28 |
| Mean/range | 13.28 | range:100-3000 | 11.44 | 83.83 | 0.2727 | 3.38 |

PIC: polymorphism information content

### Genetic diversity, genetic differentiation and gene flow in Plantago species

The highest correlations were identified between marker index with number of polymorphic band and PIC (Suppl. Table 2). The highest polymorphism was obtained in *P. amplexicaulis* (13.0%) followed by *P. pysillum* (10.1%) (Table 5). Accessions belonging to *P. lagopus* represented the lowest polymorphism (4.52%) for ISSR markers. The mean for Nei’s gene diversity (*h*) and Shannon’s information index (*I*) were 0.029 and 0.043, respectively. Accessions in *P. amplexicaulis* species represented the highest Nei’s gene diversity whilst the lowest obtained for *P. major* and *P. lanceolata* and *P. officinalis*. The results for Shannon’s information index indicated that *P. amplexicaulis* had higher variation than other species tested.

**Table 5.**
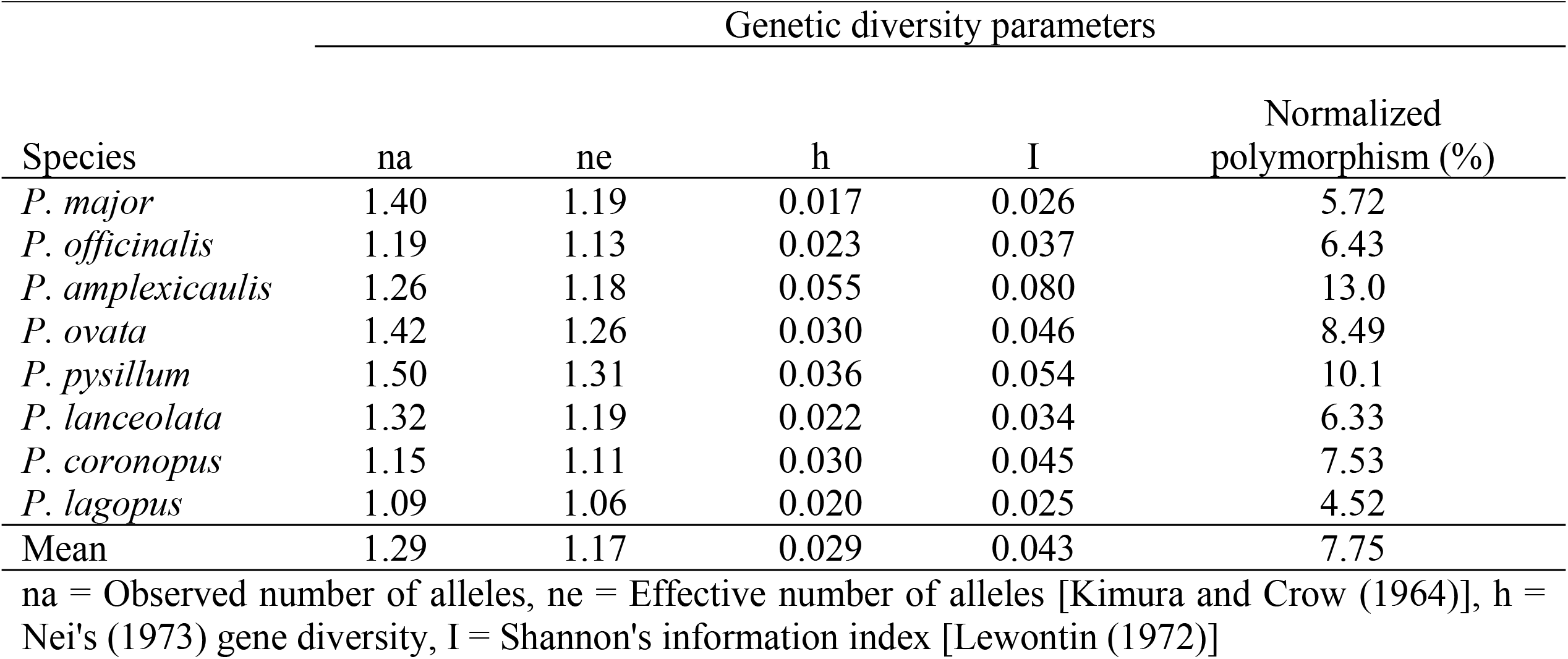
Genetic diversity parameters estimated in 31 *Plantago* accessions belonging to eight species

### Phylogenetic network of Plantago accessions

The phylogenetic network of *Plantago* accessions generated by the Neighbor-Net Algorithm is presented in Figure 4. As shown the results of Neighbor-Net analysis represented moderate split of the species tested. The output detected all the species and the network depicted moderate conflict. Of the accessions tested, two *P. amplexicaulis* samples showed conflict with long distance in the split tree diagram. Seven accessions belonging to *P. major* that were collected from various geographical regions grouped in the same group. Three accessions of *P. officinalis* constituted a distinct group at the proximity of the *P. major*. Five accessions of *P. ovata* and five belonging to *P. pysillum* grouped in the same cluster. Six accessions belonging to *P. lanceolata*, one *P. coronopus*, and two *P. lagopus* accessions constituted a separate group. Two accessions were not assigned to any cluster. The biplot of PCoA demonstrated that most of accessions belonging to the same species were clustered in same groups (Fig. 5).

**Figure 4.**
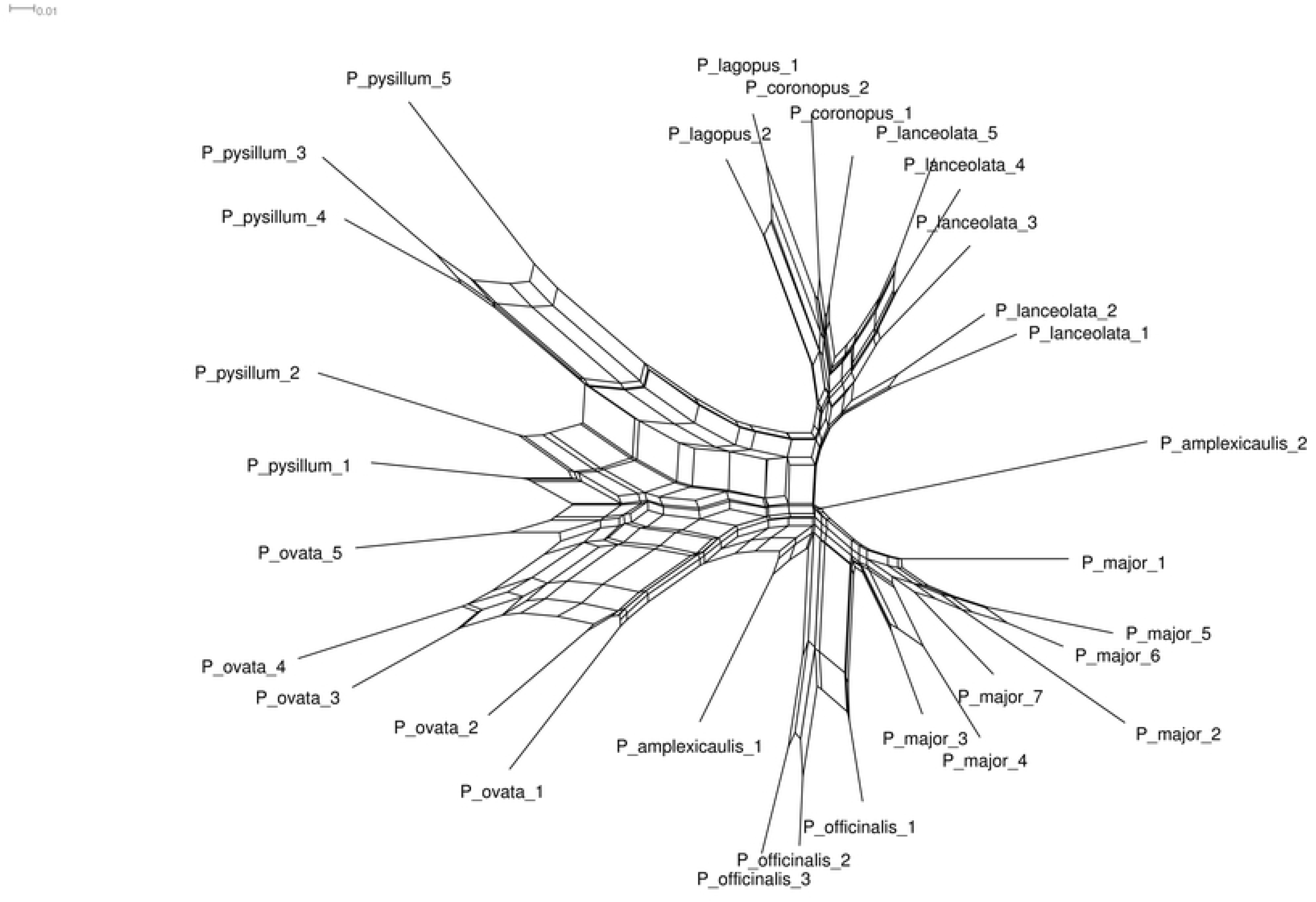
The phylogenetic network of *Plantago* accessions generated by the Neighbor-Net Algorithm

**Figure 5.**
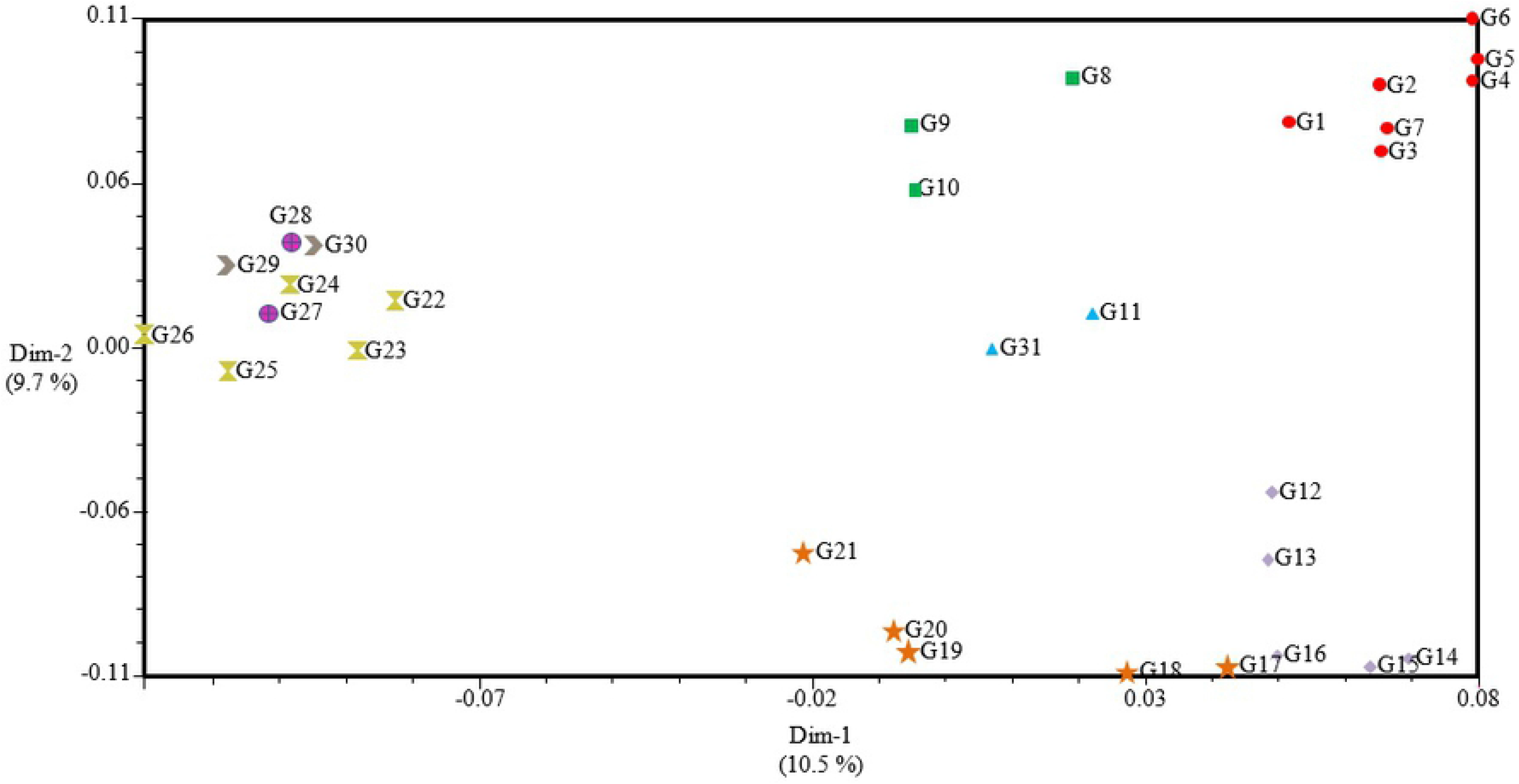
The biplot of principal coordinates analysis (PCoA) for *Plantago* accessions collected from Iran environments. Accessions belonging to same species are represented by similar symbol.

### MTAs for seed and mucilage traits

In the Manhattan plots, the highly significant MTAs were identified for each of the tested traits (Fig. 6). Fifty-six significant MTAs were detected for the traits tested in *Plantago* accessions, of which six (*H-UBC807*, *I-UBC807*, *O-UBC836*, *I-ISSR8565*, *I-IS01*, *I-ISSR8*) were shared between three seed and mucilage traits and 24 were common between two traits (Table 6). Other MTAs were trait-specific. The highest common MTAs were found for seed and mucilage yields whilst mucilage content showed less MTAs being common with the remainders of traits. Among the trait specific markers, the highest number linked markers were identified for mucilage content. The coefficient of determination of MTAs ranged from 32% for *C-ISSR2* marker linked to seed and mucilage yields to 73% for *H-UBC807* associated with seed yield.

**Figure 6.**
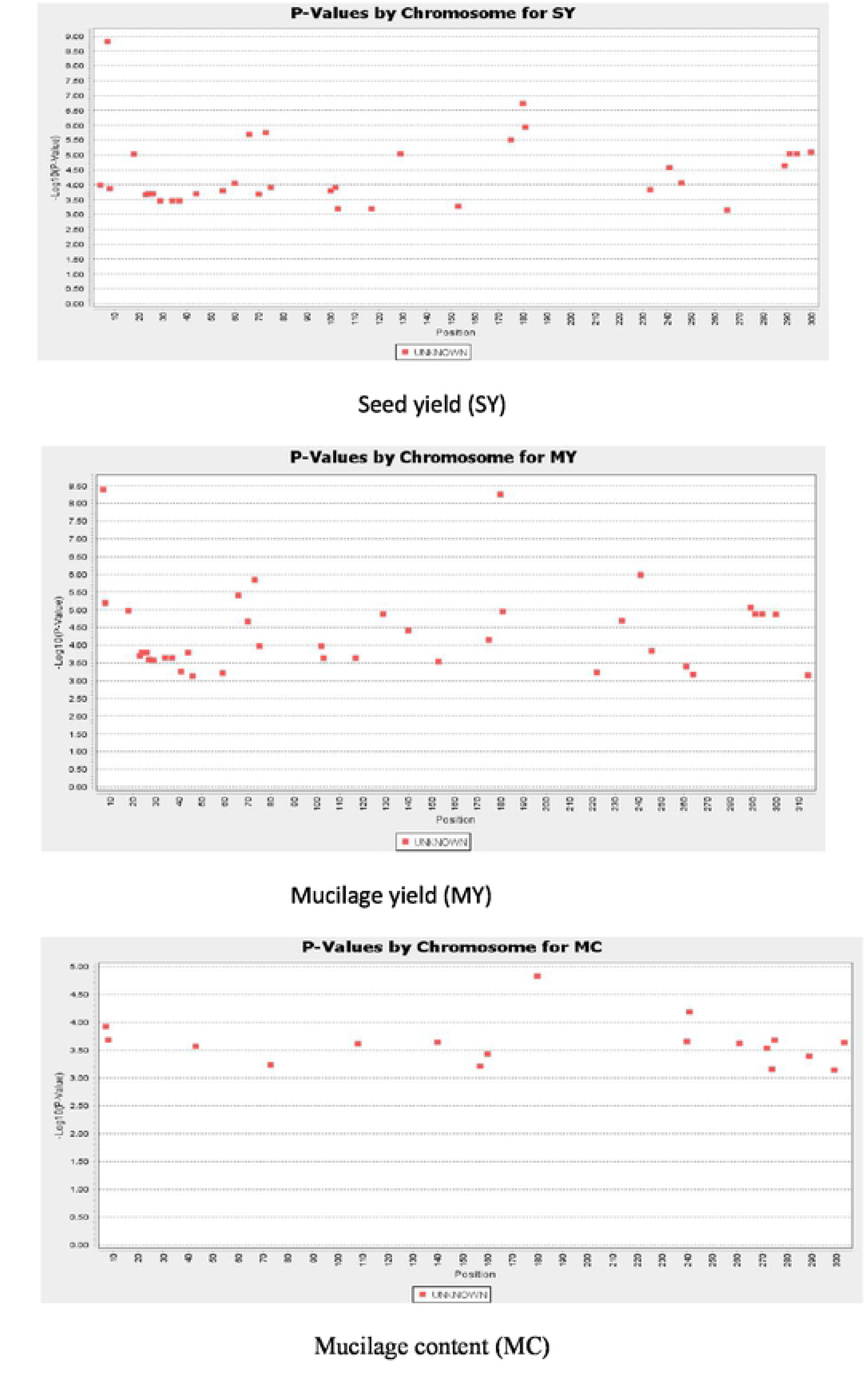
Manhattan plot displaying ISSR marker-trait associations (MTAs) identified for seed and mucilage yield in Plantago accessions belonging to eight species. Redline is significance threshold of 5% Bonferroni correction while blue line is significant threshold of 5% false discovery rate (FDR). The Y axis is the chromosome number where, 1=1A, 2=1B, 3=1D, 4=2A, 5=2B, 6=2D, 7=3A, 8=3B, 9=3D, 10=4A, 11=4B, 12=4D, 13=5A, 14=5B, 15=5D, 16=6A, 17=6B, 18=6D, 19=7A, 20=7B, 21=7D, 22= unknown chromosome

**Table 6.** Linked inter simple sequence repeats (ISSR) markers identified for seed and mucilage related traits in *Plantago* accessions tested under field conditions.

| Between –trait common marker | R<sup>2</sup> for Common markers |  |  | Trait-specific marker | R<sup>2</sup> for Trait-specific markers |  |  |
| --- | --- | --- | --- | --- | --- | --- | --- |
|  | Seed yield | Mucilage yield | Mucilage content |  | Seed yield | Mucilage yield | Mucilage content |
| *H-UBC807* | 0.73 | 0.72 | 0.42 | *E-UBC807* | 0.42 | - | - |
| *I-UBC807* | 0.41 | 0.52 | 0.39 | *F-UBC835* | 0.4 | - | - |
| *D-UBC811* | 0.51 | 0.51 | - | *B-UBC880* | 0.4 | - | - |
| *I-UBC811* | 0.39 | 0.40 | - | *E-ISSR6* | 0.34 | - | - |
| *J-UBC811* | 0.39 | 0.40 | - | *B-UBC836* | 0.43 | - | - |
| *K-UBC811* | 0.39 | 0.40 | - | *M-UBC811* | - | 0.38 | - |
| *L-UBC811* | 0.39 | 0.40 | - | *I-UBC825* | - | 0.35 | - |
| *O-UBC811* | 0.37 | 0.38 | - | *N-UBC825* | - | 0.34 | - |
| *B-UBC825* | 0.37 | 0.39 | - | *A-UBC836* | - | 0.35 | - |
| *E-UBC825* | 0.37 | 0.39 | - | *H-ISSR15* | - | 0.46 | - |
| *L-UBC825* | 0.39 | 0.40 | - | *G-IS16* | - | 0.35 | - |
| *H-UBC836* | 0.56 | 0.54 | - | *A-ISSR6* | - | 0.37 | - |
| *L-UBC836* | 0.39 | 0.40 | - | *D-ISSR6* | - | 0.34 | - |
| *O-UBC836* | 0.56 | 0.57 | 0.35 | *S-ISSR9* | - | 0.34 | - |
| *Q-UBC836* | 0.42 | 0.42 | - | *K-UBC825* | - | - | 0.38 |
| *F-UBC880* | 0.42 | 0.42 | - | *D-UBC889* | - | - | 0.39 |
| *G-UBC880* | 0.35 | 0.39 | - | *H-ISSR15* | - | - | 0.39 |
| *C-ISSR2* | 0.35 | 0.32 | - | *G-ISSR20* | - | - | 0.35 |
| *I-ISSR3* | 0.51 | 0.50 | - | *J-ISSR20* | - | - | 0.37 |
| *C-ISSR20* | 0.35 | 0.38 | - | *H-IS01* | - | - | 0.39 |
| *D-ISSR8565* | 0.55 | 0.44 | - | *A-ISSR6* | - | - | 0.39 |
| *I-ISSR8565* | 0.63 | 0.44 | 0.49 | *L-ISSR6* | - | - | 0.38 |
| *J-ISSR8565* | 0.58 | 0.71 | - | *N-ISSR6* | - | - | 0.34 |
| *A-IS01* | 0.41 | 0.48 | - | *O-ISSR6* | - | - | 0.39 |
| *I-IS01* | 0.47 | 0.58 | 0.44 | *D-ISSR9* | - | - | 0.34 |
| *N-IS01* | 0.43 | 0.41 | - | *H-ISSR9* | - | - | 0.39 |
| *I-ISSR8* | 0.48 | 0.51 | 0.36 |  |  |  |  |
| *K-ISSR8* | 0.51 | 0.5 | - |  |  |  |  |
| *N-ISSR8* | 0.51 | 0.5 | - |  |  |  |  |
| *E-ISSR9* | 0.52 | 0.5 | - |  |  |  |  |

\*R<sup>2</sup>: coefficient of determination showing contribution of each ISSR marker

## Discussion

The application of molecular markers has a vital role in genetic diversity analysis of biological samples. In the present study, genetic diversity of several *Plantago* species was assessed using ISSR technique. Estimation of genetic diversity of plant materials is one of the most important pre- breeding activities that assist conservation of plant species. The results of ISSR analysis in the present study demonstrated genetic diversity in *Plantago* species collected from various regions of Iran environments. In a study with *Plantago brutia*, the results of ISSR analysis demonstrated high genetic diversity at the species level [33]. Several ISSR primers tested in our study yielded 100% polymorphism in the tested *Plantago* species. In the Osmanaand Abdein [1]) study, one of ISSR primers presented 80% polymorphism among six *Plantago* species whilst several RAPD and SCOT markers showed 100% polymorphism. Although RAPD markers yield high polymorphism in biological samples, ISSR is more reliable technique for genetic mapping and diversity analysis [34, 35]. Presently, analysis of 25 ISSR primers resulted in above 80% polymorphism detected in the 31 *Plantago* accessions. In other studies with *Plantago* 19.35% (Kaswan et al. [4] and 70.54% [36] polymorphisms were identified for ISSR primers tested in *P. ovata* and *P. psyllium*, respectively. Analysis of PIC analysis provides an estimate of the discriminating power of a marker. The mean PIC obtained for the ISSR primers (=0. 2727) in the present study was lower than the mean PIC (=0.36) estimated for 12 ISSR primers in *P. psyllium* in the Rahimi et al. [36] study. This result shows the ISSR primers tested in our study were informative with respect to discerning genetic diversity of the eight *Plantago* species. The marker index estimated in the present study demonstrated the potential of the ISSR primers for detection of polymorphism among *Plantago* species at DNA level. Analysis of several DNA markers in *Musa acuminatacolla* demonstrated that ISSR resolved highest marker index (16.39) compared with RAPD and SPAR markers [37].

Genetic diversity of *Plantago* accessions belonging to the eight species were also tested using Nei’s gene diversity (*h*) and Shannon’s information index (*I*)*. P. amplexicaulis* and *P. psyllium* showed highest polymorphism for ISSR markers. One of reasons for higher polymorphism in these two species could be their mating system; one is inbreeder and the other shows a mixture of open and selfed pollinations behavior. Basically, higher genetic diversity in populations could be due to mating systems or habitat/dynamics history [33,38]. Self-incompatibility is another reason for genetic diversity that has been demonstrated in several *Plantago* spp. (*P. brutita*, *P. media*). The results of both the Nei and Shannon indices were consistent for the *P. amplexicaulis* species with the highest genetic diversity identified in our study. For other species the results of the two indices were almost similar. Estimation of these indices demonstrated higher genetic diversity for accessions in *P. amplexicaulis* and *P. psyllium* compared with other *Plantago* species tested.

Marker-trait linkages help identify MTAs and the linked markers could be used for marker-assisted selection (MAS) programs in breeding for a target trait in plant species. The results of the present study showed that several ISSR markers were linked to multiple traits in *Plantago*. In a study with agronomic trait in cumin (*Cuminum cyminum*), five markers were associated with more than one trait [39]. So far, only one study [40] has been reported for analysis of the association of agronomic traits and DNA markers in *Plantago* but number of markers (24 SCoT), species (only two) and tested environments were rather lower than those tested in the present study. Accordingly, the results of the present study offer a valuable first insight into genetic control of mucilage and seed yield traits for use in MAS in *Plantago*. For MAS, the marker should be highly polymorphic and should discriminate between different genotypes in breeding material [41]. The level of polymorphism in ISSR markers tested in the present study was rather high being applicable for use in MAS for breeding *Plantago*. In addition, the results of this study will help *Plantago* breeders to select accessions for a collection of *Plantago* species in the germplasm bank of medicinal plants.

## Conclusions

Further insights regarding phylogenetic network of *Plantago* individuals collected from various geographical regions of Iran were revealed by the ISSR marker data. The ISSR data and genetic variation parameters demonstrated large polymorphism was exist among accessions of the eight *Plantago* species. The results of Neighbor-Net analysis demonstrated moderate split and relatively low conflict for species discrimination. Several ISSR-trait associations were identified for seed yield and mucilage traits of which those that were shared between tested traits are good candidate for use in MAS programs. The *Plantago* groups were notably highly correlated to the taxonomic status of the *Plantago* samples tested. In conclusion, information on the genetic diversity of the selected accessions is of ultimate importance as it contributes to the information on the species and allows accession selection for future *Plantago* improvement and conservation programs.

## Availability of data and materials

All data and materials used in this work were publicly available.

## Ethics approval and consent to participate

The ethical approval or individual consent was not applicable

## Authors’ contributions

MT contributed to study design, the literature search, data collection, data analysis, software working. BH defined and supervised the project and wrote the first and final draft of the manuscript. ZS collected the data for mucilage. AD helped technical works and edited the final draft of the manuscript. ZH helped in figure construction and data analysis. CMR involved in data analysis and editing the first and final draft of the manuscript

## Consent for publication

Not applicable.

## References

1. Osman AKE, Abedin MAE.Karyological and molecular studies between six species of *Plantago* in the Northern border region at Saudi Arabia. J Taibah Univ. Sci. 2019; 13(1): 297–308.

2. Shahriari Z, Heidari B, Dadkhodaie A, Richards, CM. Analysis of karyotype, chromosome characteristics, variation in mucilage content and grain yield traits in *Plantago ovata* and *P. psyllium* species. Ind. Crops Prod. 2018; 123: 676–686.

3. Shahriari Z,. Heidari B, Dadkhodaie, A. Dissection of genotype × environment interactions for mucilage and seed yield in Plantago species: Application of AMMI and GGE biplot analyses. PLoS ONE 2018; 13(5), e0196095. https://doi.org/10.1371/journal.pone.0196095

4. Kaswan V, Joshi AJ, Malo, SR. Assessment of genetic diversity in Isabgol (*Plantago ovata* Forsk.) using random amplified polymorphic DNA (RAPD) and inter-simple sequence repeat (ISSR) markers for developing crop improvement strategies. Afric. J Biotechnol. 2013; 12(23): 3622–3635.

5. Mirmasumi M, Ebrahimzadeh H, Tabatabaei SMF. Mucilage production in tissue culture of *Plantago lanceolata*. J. Agric. Sci. Technol. 2001; 3: 155–160.

6. Haddadian K, Haddadian K, Zahmatkash, M. A review of Plantago plant. Int J Trad Knowl. 13 (4) 6: 81–685.

7. McNeely JA, Miller KR, Reid WV, et, al. Conserving the world’s biological diversity. World Conservation Union, World Resources Institute, World Wildlife Fund–US, and the World Bank.1990.

8. Vicente, M.J., Segura, F., Aguado, M., Migliaro, D., Franco, J.A., Martínez-Sánchez, J.J., 2011. Genetic diversity of Astragalus nitidiflorus, a critically endangered endemic of SE Spain, and implications for its conservation. Biochem. Syst. Ecol. 39, 175–189.

9. Ferreira V, Gonçalves S, Matos M, Correia S, Martins N, Romano, A. Pinto-Carnide O. Genetic diversity of two endemic and endangered *Plantago* species. Biochem. Syst. Ecol. 2013; 51: 37–44.

10. Sharma K, Agrawal V, Gupta S, Kumar R, Prasad, M. ISSR marker-assisted selection of male and female plants in a promising dioecious crop: jojoba (*Simmondsia chinensis*). Plant Biotechnol. Rep. 2008; 2: 239–243

11. Xue-En F, Qin C, Li-Ping Y, Wei, W. Application of ISSR in genetic relationship analysis of sorghum species. Acta Agron. Sinic. 2008; 34(8): 1480–1483.

12. Godvin ID, Aikten EAB, Smith, LW. Application of inter simple sequence repeat (ISSR) markers to plant genetics. Electrophoresis, 1997; 18: 1524–1528.

13. Menon PS. ISSR scientists develop new clove varieties. J Herbs Spice Medicin Plants 2000; 7(1): 103–105.

14. Lu Z, Duan H, Lu L, Sun X.Genetic relationships of *Osmanthus* based on ISSR-PCR. Biologia, 2010; 65(3): 459–464.

15. Bamhania K, Khatakar S, Punia A, Yadav, OD. Genetic variability analysis using ISSR markers in *Withania Somnifera* L. dunal genotypes from different regions. J Herbs Spices Medicine Plants 2013; 19: 22–32.

16. Seyedimoradi H, Talebi R.Detecting DNA polymorphism and genetic diversity in Lentil (*Lens culinaris* Medik.) germplasm: comparison of ISSR and DAMD marker. Physiol. Mol. Biol. Plants 2014; 20(4): 495–500. DOI 10.1007/s12298-014-0253-3.

17. Rajendran HAD, Muthusamy R, Stanislaus AC, Krishnaraj T, Kuppusamy T, Ignacimuthu S, Al-Dhabi NA. Analysis of molecular variance and population structure in southern Indian finger millet genotypes using three different molecular markers. J Crop Sci. Biotechnol. 2016; 19 (4): 275–283.

18. De Vita A, Bernardo L, Gargano D, Palermo AM, Mussachio A.Investigating genetic diversity and habitat dynamics in *Plantago brutia* (Plantaginaceae), implications for the management of narrow endemics in Mediterranean mountain pastures. Plant Biol. 2009; 11: 821–828.

19. Murray MG, Thompson, WF. Rapid isolation of high molecular weight plant DNA. Nucl. Acids Res. 1980; 8(19): 4321–4326.

20. Yeh FC, Yang R, Boyle T, Ye ZH, Mao JX.POPGEN Ver. 1.32. The user-friendly software for population genetic analysis. 1997, Molecular Biology and Biotechnology Center, University of Alberta, Alberta, Canada.

21. Shannon CE. A mathematical theory of communication. The Bell System Technical Journal, 1948: 27: 379–423 and 623–656.

22. Anderson A, Churchill GA, Autrique JE, Tanksley SD, Sorrells, ME. Optimizing parental selection for genetic linkage maps. Genome, 1993: 36(1): 181–186.

23. Kumar A, Mishra P, Singh SC, Sundaresan, V. Efficiency of ISSR and RAPD markers in genetic divergence analysis and conservation management of *Justicia adhatoda* L., a medicinal plant. Plant Syst. Evol. 2014; 300: 1409–1420.

24. Nei M. Genetic distance between populations. The American Naturalist, 1972: 106(949): 283–292.

25. Nei M.Analysis of gene diversity in subdivided populations. Proceedings of the National Academy of Sciences, 1973; 70(12): 3321–3323.

26. Leberg PL. Estimation of allelic richness: effects of sample size and bottlenecks. Mol. Ecol. 2002; 11: 2445–2449.

27. Bryant D, Moulton, V. Neighbor-net: An agglomerative method for the construction of phylogenetic networks. Mol. Biol. Evol. 2004; 21: 255–265.

28. Huson DH, Bryant, D. Application of phylogenetic networks in evolutionary studies. Mol. Biol. Evol. 2006; 23: 254–267.

29. Bradbury PJ, Zhang Z, Kroon DE, Casstevens TM, Ramdoss Y, Buckler, ES. TASSEL: software for association mapping of complex traits in diverse samples. Bioinformatics 2007; 23: 2633–2635. https://doi.org/10.1093/bioinformatics/btm308

30. Yu J, Pressoir G, Briggs WH, Vroh Bi I, Yamasaki M, Doebley, JF. et al. A unified mixed-model method for association mapping that accounts for multiple levels of relatedness. Nat. Genet. 2006; 38(2): 203–208 doi:10.1038/ng1702.

31. Myles S, Peiffer J, Brown PJ, Ersoz E, Zhang Z, Costich DE, Buckler, ES. Association mapping: Critical considerations shift from genotyping to experimental design. Plant Cell 2009; 21: 2194–2202. Available online at: http://dx.doi.org/10.1105/tpc.109.068437

32. Zhu C, Gore M, Buckler ES, Yu J Status and prospects of association mapping in plants. Plant Genome 2008; 1:5–20. Available online at: http://dx.doi.org/10.3835/plantgenome2008.02.0089

33. Vita DA, Bernardo L, Gargano D, Palermo AM, Peruzzi L, Musacchio, A. Investigating genetic diversity and habitat dynamics in Plantago brutia (Plantaginaceae), implications for the management of narrow endemics in Mediterranean mountain pastures. Plant Biology. 2009; 11: 821–828.

34. Chen L, Chen F, He S, Ma, L. High genetic diversity and small genetic variation among populations of *Magnolia wufengensis* (Magnoliaceae), revealed by ISSR and SRAP markers. Elec. J Biotechnol. 2014; 17, 268–274.

35. Sarwat M. ISSR: a reliable and cost-effective technique for detection of DNA polymorphism. Methods Mol. Biol. 2012; 862: 103–21. doi: 10.1007/978-1-61779-609-8_9.

36. Rahimi M, Hatami Malek H, Mortezavi, M. Identification of informative markers of agronomic traits in different ecotypes of sand plantain (*Plantago psyllium*). Biologia, 2017; 63 (4): 325–333.

37. Lamare A, Rao, SR. Efficacy of RAPD, ISSR and DAMD markers in assessment of genetic variability and population structure of wild *Musa acuminatacolla*. Physiol. Mol. Biol. Plants 2015; 21(3), 349–358.

38. Schaal BA, Hayworth DA, Olsen KM, Rauscher JT, Smith, WA. Phylogeographic studies in plants: problems and prospects. Mol. Ecol. 1998; 7: 465–474.

39. Archangi A, Heidari D, Mohammadi-nejad G. Association between seed yield-related traits and cDNA-AFLP markers in cumin (*Cuminum cyminum*) under drought and irrigation regimes. Ind Crop Prod. 2019; 276–283.

40. Rahimi M, Nazari L, Kordrostami M, Safari, P. SCoT marker diversity among Iranian Plantago ecotypes and their possible association with agronomic traits. Scientia Horticulturae 2018; 233: 302–309.

41. Collard CVB, MacKill, D. Marker-assisted selection: an approach for precision plant breeding in the twenty-first century. Phil. Trans. R. Soc. B. 2008; 363: 557–572.

